# Forest edge effects on termites in a neotropical rainforest

**DOI:** 10.64898/2026.10.28.762416

**Authors:** Edouard Duquesne, Antoine Dumortier, Camille Legrand, Pavel Babczenko, Yves Roisin

## Abstract

Termites are dominant arthropods in tropical forests, where they play a major role in organic matter decomposition. However, their activity may be reduced by habitat degradation, to which soil-dwelling species are especially sensitive. Nowadays, one of the major threats affecting tropical forests is fragmentation, through two concurring effects: a reduction in area of the forested zones, and an increase in the length of forest edges. These effects are hard to disentangle. In this study, we took advantage of a powerline clearing to evaluate edge effects on the termite fauna of an otherwise continuous rainforest in French Guiana. No edge effect was detected 100–300 m into the interior. Soil feeders were not significantly less present at the forest edge although they have an allegedly thinner skeleton and a need for moist and soft soil. Finally, no increase in beta diversity near the edge was found either. Overall, it seems that termites are resilient insects against forest edges as long as the fragment size stays large enough. However, a 3-fold larger presence of an opportunistic and pest species (*Heterotermes tenuis*) was found at the forest edge. We hypothesize that *H. tenuis* good adaptability and dispersibility may have allowed it to quickly colonize and dominate this new environment before more cryptic species, as it’s been shown for ants.

## 1 INTRODUCTION

In the Anthropocene, biodiversity has faced multiple pressures such as land use, pollution, climate change, and invasive species (Pereira *et al*., 2012). As a result, anthropogenic biomes now make up more than 75% of all ice-free land on Earth (Ellis & Ramankutty, 2008). Moreover, land use and fragmentation have increasing impacts on biodiversity as climate change creates a snowball effect (Mantyka-Pringle *et al*., 2012; Seidl *et al*., 2011). Forests are heavily affected by deforestation, especially in the tropics for meat and oleaginous production (Marques *et al*., 2019). The resulting fragmentation is such that more than 70% of worldwide forests are within 1 km of a forest edge (Haddad *et al*., 2015). Even worse, the Brazilian Atlantic Forest is now so fragmented that almost 50% of the remaining forested area is within 100 m from an open habitat, and more than 70% is situated at less than 250 m from the edge (Ribeiro *et al*., 2009).

Fragmentation is a division process which involves two concurring effects: (1) habitat area reduction and (2) edge length increase (Fahrig, 2003; Hadley & Betts, 2016). Both may affect flora and fauna in general or specific ways: (1) fragments of reduced areas may be too small to provide enough resources to sustain a viable population; even in larger fragments, local populations may fail to harbor a sufficient genetic diversity to survive in the long run; (2) forest edges experience abiotic and biotic conditions very different from the forest core, and as the ratio *edge length/core area* increases, communities will be increasingly affected (Cordeiro *et al*., 2015; Harper *et al*., 2005; Laurance *et al*., 2002). Unfortunately, the effects of area reduction and forest edges are hard to study independently (Fahrig, 2017).

Newly generated forest edges are usually characterized by broader temperature fluctuations, a higher luminosity and stronger winds compared to forest cores (Laurance, 2004). The matrix surrounding the forest fragments also offers possibilities of colonization and crossing for more opportunistic species, and as a result invasive species are typically gaining more from forest edges than endemic species (Pfeifer *et al*., 2017). All those changes modify species composition due to new biotic and abiotic interactions as well as differences in physiological resistance (Murcia, 1995). In this manner, more than 85% of vertebrate species abundance are influenced by forest edges either positively or negatively, depending on their thermal regulation, their surface/volume ratio and their travel capacity (Pfeifer *et al*., 2017). Although arthropods represent more than 42% of the animal biomass (Bar-On *et al*., 2018), most studies on diversity loss focus on plants and vertebrates (Ceballos *et al*., 2015). However, arthropods and especially insects are suffering from massive depletion (Hallmann *et al*., 2017; Seibold *et al*., 2019) and this will only get worse as climate change increases (Halsch *et al*., 2021).

The reduction of suitable habitats has been studied for a long time and its consequences on biodiversity are well known. On the other hand, forest edges seem to have different outcomes depending on the species. For insects, it seems that forest edges are mainly negative for social and soil-feeding species but mainly positive for invasive and flying species (Caitano *et al*., 2020). Microclimatic effects created by an edge will typically die out within 100 m from the edge (Dambros *et al*., 2013; Laurance *et al*., 2002) but this distance can increase during drier periods (Kunert *et al*., 2015). However, its impacts on insect communities can go deeper in the forest, as much as 1 km for dung beetles in a New Zealand forest (Ewers & Didham, 2008) and 200 m for Amazonian leaf-litter ants (Carvalho & Vasconcelos, 1999)

Termites (Blattodea, Isoptera) are mainly present in tropical and subtropical regions (Eggleton *et al*., 1994). As of 2013, more than 3106 species and 331 genera have been described but new species are being discovered every year (Krishna *et al*., 2013). They are among the most abundant insects in tropical rainforests where they play a major role as decomposers and ecosystem engineers (Holt & Lepage, 2000; Jouquet *et al*., 2016). They feed on organic matter in decomposition such as dead wood, leaves, roots, feces, soil, and humus (Eggleton & Tayasu, 2001; Freymann *et al*., 2008). Typically, two guilds are differentiated: soil feeders and wood feeders. Soil feeders, through their food regime and digestive system, play a major role in the mineralization of soil nitrogen (Ji & Brune, 2006). Wood feeders are notably responsible for the recycling and breaking of one-third of the annual production of dead wood (Verma *et al*., 2009). The sensitiveness of termites to disturbance, including man-generated edge effects, varies between guilds: in African rainforests, soil feeders were shown to suffer more than wood feeders from anthropogenic disturbances (Eggleton *et al*., 2002). Wood feeders seem to be more resistant to drought than soil feeders because the latter have a thinner exoskeleton and need wet, soft soils (Davies *et al*., 2012; Eggleton *et al*., 2002; Muvengwi *et al*., 2017). Because the species-richest and most specialized termite family (Termitidae) diversified in the early Cenozoic in Africa and most species are poor dispersers, termite phylogenetic lineages follow strong zoogeographic patterns and generate very distinct assemblages on different continents (Bourguignon *et al*., 2017). The response of neotropical termite communities to edge effects may therefore differ from those of Afrotropical ones.

In neotropical forests, edge effects on termites have been studied by Dambros *et al*. (2013) and Almeida *et al*. (2017) in Brazil. However, in both studies, the forest fragments were too small to truly discriminate edge effects from those of habitat reduction. Both studies found that species composition differed between the forest edge and the core. Dambros *et al*. (2013) found that a road-associated edge had an impact on soil moisture up to 81 m into the forest, but the sampled forest was itself engulfed in a large urban agglomeration, which may have produced confounding effects. In French Guiana, the construction of the dam and hydroelectric plant of Petit Saut in the 1990s required clearings to be made in the forest, first for the access road, second for the power lines carrying electricity from the dam to the coastal areas. The powerline cut consists in a series of straight segments, about 90 m wide, regularly cleared from shrubs and arborescent vegetation. This setting provides the opportunity to study edge effects in a forest that is otherwise continuous and whose fauna is not constrained by fragment area, and allowed us to test the following predictions.

(1) Wood-feeding termites are allegedly more resistant to dry conditions than soil-feeding termites because the latter have a thinner exoskeleton and a food regime requiring a soft and moist soil (Davies *et al*., 2012; Eggleton *et al*., 2002; Muvengwi *et al*., 2017). Therefore, wood feeders should be less affected by climatic fluctuations near the forest edge and could even benefit from an increased amount of litter and dead wood.

(2) Beta-diversity may increase along the edge, if disturbance favors opportunistic species with a high capacity of dispersion and colonization (Ivanov & Keiper, 2010; Pfeifer *et al*., 2017), which enhance the turnover of species between sites (Krishnadas *et al*., 2019; Swart *et al*., 2019).

(3) Considering that microclimatic effects typically die out within 100 m of an edge (Ewers & Didham, 2008), and that other litter-dwelling invertebrate communities can be affected as far as 200–300 m into the interior of the forest (Carvalho & Vasconcelos, 1999; Marsh *et al*., 2018), we expected the edge effects on termite communities to reach about the same depth.

## 2 METHODS

### 2.1 Study sites

The study took place in French Guiana, in the Amazonian forest along the road connecting the hydroelectric dam of Petit Saut to the coastal road (N1) between Kourou and Sinnamary. Three sites (L, N and M, coordinates below) were sampled in 2019, 2020 and in 2022 during the small dry season from February to March. All the sites were situated next to a relatively straight edge 90 m wide made for the powerline carrying electricity from the Petit Saut dam to coastal areas. The three sites were all separated by at least 3000 m (Figure 1). The studied area is characterized by hills and plateaus within an important hydrographic network and with mainly plants of the Lecythidaceae and Caesalpinioideae (sub)families (Guitet *et al*., 2015). The dominant soil is acrisol, with an important level of clay and a yellow-red color due to iron oxides (Barret & Vandé, 2001; Guitet *et al*., 2015). The forest experiences on average 2750 mm of rain per year. However, there is a strong seasonality: during the rainy season (April to July), rainfall may add up to 350 mm per month while it drops to 50 mm per month during the dry season (July to November). Annual temperature is on average 26°C (Barret & Vandé, 2001).

**Figure 1.**
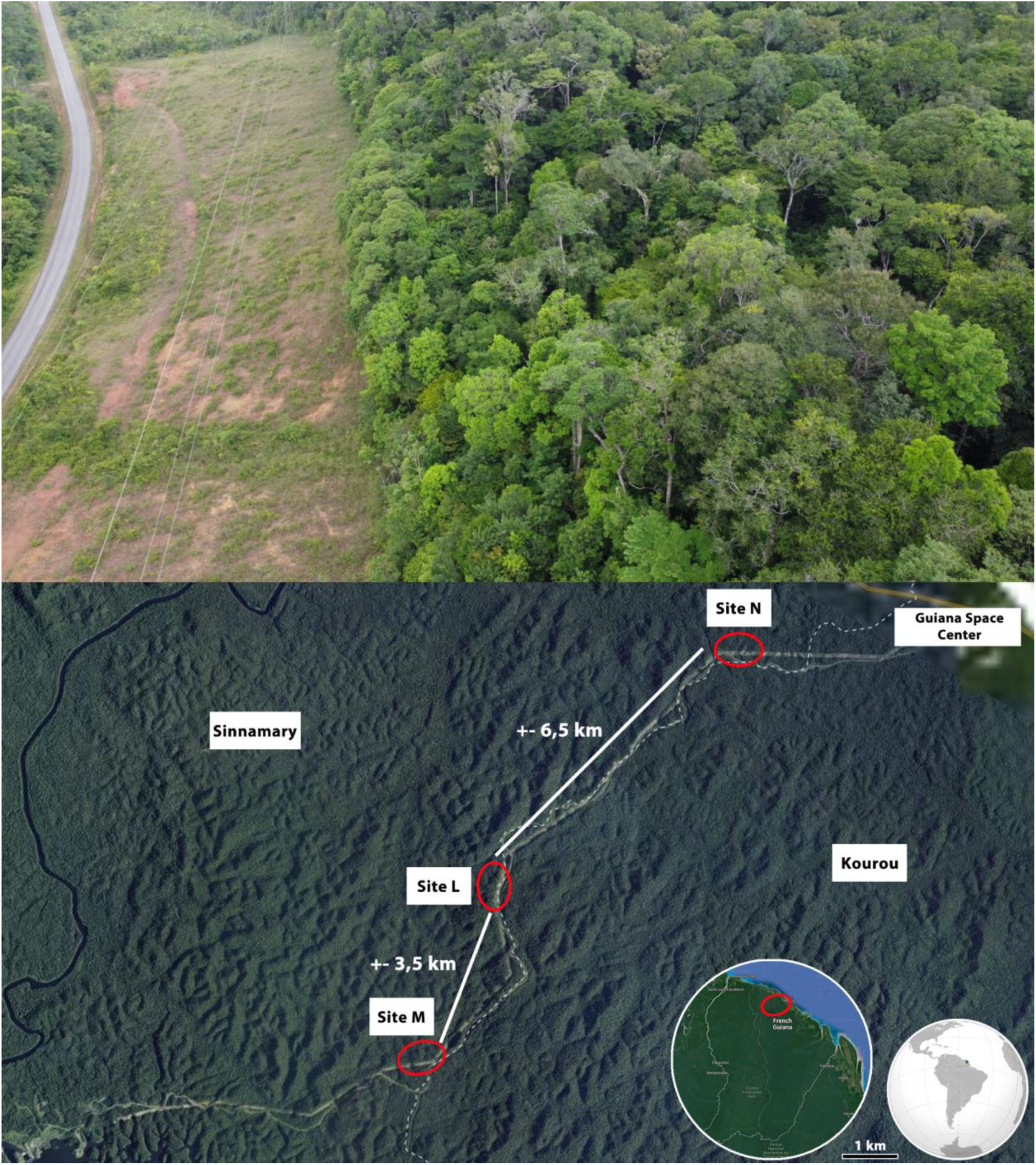
Top: aerial picture of the forest edge at the site L. The paved road on the left is 6.2 m wide. Bottom: location of the three sites studied in the Sinnamary commune of French Guiana. Satellite images from IGN and TomTom (2022) and South America map from Wikipedia.

### 2.2 Termite sampling

Termites were collected using the Jones & Eggleton (2000) protocol slightly modified as recommended by Roisin & Leponce (2004). The quadrats and transects of the three sites (**L** = 5°06’41“N 52°57’55”W, **N** = 5°09’02.4“N 52°55’33.7”W and **M** = 5°04’51“N 52°58’41”W) were positioned using a compass, a decameter and a GPS (Garmin eTrex 32). For the edge effect distance (1), six transects (240 m long) were made parallel to the forest edge of the L site at 10 m, 30 m, 60 m, 100 m, 250 m, and 500 m from the cut. For the second part, transects were made at 10 m and 100 m (except for the N21 quadrat which was made at 90 m from the forest edge due to the presence of a big treefall) in the N and M sites. Each transect was made out of 25 quadrats of 5 m^2^ (2.24 × 2.24 m), each spaced by 10 m, center to center. Each quadrat was delimited using a rope and tent pegs. Inside each quadrat, dead wood, nests, litters, and trunks as well as six soil blocks of 12 × 12 × 10 cm were carefully searched by two persons for around 15 min in order to collect a dozen individuals from each termite colony or foraging party. The termites were then conserved in 100% ethanol for morphological and molecular analyses and/or in 80% ethanol for anatomical analyses. In total, 250 quadrats were sampled over the three years.

### 2.3 Termite identification

First, samples were classified according to the morphology of the workers and soldiers and identified to the genus using a dissecting microscope (Leica MZ06) and Constantino’s (2002) key. Various keys were then used to identify the species, depending on the genus (Emerson, 1925; Mathews, 1977). For soldierless species, the enteric valve situated at the junction of the first (tubular) section of the hind gut with the paunch was dissected, splained and mounted on a glass slide in a droplet of polyvinyl alcohol-based mounting fluid (CMCP-9, Polysciences Inc.). Specimens were assigned to (morpho)species according to their gut anatomy and enteric valve armature, by comparison with published data on neotropical Apicotermitinae and our own database, assembled from Bourguignon *et al*. (2010, 2013, 2016). Samples proving more difficult were identified through sequencing. For this part, the cytochrome c oxidase subunit 2 (COII) was chosen for its variability, small size (between 700 and 1000 bp) and broad sequence availability for termites (Huang *et al*., 2011; Ohkuma *et al*., 2004). The termites’ DNA was extracted from their head (to reduce any gut contamination) using NucleoSpin®Tissue (Macherey-Nagel). DNA amplification was done using the kit BigDye V3.1 (Applied Biosystems™) with two primers ModAtLeu (CAG ATA AGT GCA TTG GAT TT, Miura *et al*., 2000) and B-tLys (GTT TAA GAG ACC AGT ACT TG, Simon *et al*., 1994). The amplification was performed on a thermocycler (Eppendorf Mastercycler x50S: 1 min at 96°C, followed by 25 cycles of: 10 s at 96°C, 5 s at 50°C and 4 min at 60°C). Once purified, the sequencing was performed on a capillary sequencer, a DNA Analyser 3730 (Applied Biosystems™). Sequences were then corrected and aligned using CodonCode Aligner (CodonCode Corporation). Finally, sequences were compared by using the BLAST tool from NCBI (2022).

### 2.4 Environmental variables

Several environmental variables were collected for the edge effect distance (1). A surface litter of 1 m^2^ was collected at each quadrat from each transect (10, 30, 60, 100, 250 and 500) at site L in 2020. The drying oven was not big enough, so the litter samples were dried off by the sun. As there was still a little bit of humidity, a small sample of 15 g from each quadrat was placed in a small drying oven in order to evaluate the degree of humidity left in the samples. The canopy openness was also evaluated in 2019 (transects 10, 30, 60 and 100) and 2020 (250 and 500) by taking hemispherical pictures which were then treated on Hemispherical_2.0 (Beckschäfer, 2015) with ImageJ software (Rasband, 1997). Deadwood volume was assessed using the method of Van Wagner (1968): a line was drawn along each transect (*L* = 6 x 240 m = 1440 m) and the diameter (*d*) of each deadwood ≥ 10 cm crossing the line up to 2 m high was taken into consideration. The full volume per air unit (m^3^/m^2^) was calculated using the following formula: *V* = (*π*^2^ × ∑ *d*^2^) × 8*L*. Finally, the elevation was measured using the GPS described earlier.

### 2.5 Data analyses

Data were analyzed using the statistical software R (Version 4.2.1; R Core Team, 2021) with several packages. Before any statistical analysis, conditions of normality were tested through a Shapiro test (Shapiro & Wilk, 1965) and homoscedasticity of variance was tested through a Bartlett test (Bartlett, 1937). First, sample effort was assessed by making sample-and occurrence-based rarefaction curves from the Vegan package (Oksanen *et al*., 2010). These analyses allow us to evaluate the sampling effort as well as interpolating the number of species based on the number of quadrats. Completeness of each transect was assessed through the iNEXT package (Hsieh *et al*., 2016) which is based on the use of Hill numbers (here q = 0) under Bernoulli’s model. They were performed for both parts (1. Edge effect distance and 2. Edge effect and beta diversity) of this research.

Concerning the first part (1), the species relative frequencies for the six transects at site L were compared using Kruskal-Wallis test (Kruskal & Wallis, 1952) followed by a post-hoc Dunn test with the Bonferroni correction (Dunnett, 1955). The same test was applied as well for each (sub-)family, for each guild and for each environmental variable (Elevation (m), Canopy openness (%) and Litter (g)) between each transect. Shannon diversity index (H, (Shannon, 1948) of each transect was calculated using the Vegan package. Following this, linear regressions were performed between H of each transect and the transect mean of each environmental variable (Elevation (m), Canopy openness (%) and Litter (g) and Dead wood (m^3^/m^2^). Finally, a NMDS (non-metric multidimensional scaling) was performed with the Vegan package using the Bray-Curtis distance (Bray & Curtis, 1957) index on the occurrences of each species per transect.

For the second part (2), distributions of the forest edge (10 m) and the interior (100 m) were compared using a two-sample Kolmogorov-Smirnov test (Massey, 1951). Then, for each (sub)-family and each guild, species relative frequencies between the forest edge and the interior were compared using a Wilcoxon test (Mann & Whitney, 1947; Wilcoxon, 1945). Chi-square tests (Pearson, 1900) were performed between each species at the forest edge and at the interior. If for one of the transect, a species had less than 5 occurrences, a Fisher test (Fisher, 1992) was performed instead, as recommended by Siegel (1957) and McCrum-Gardner (2008). A NMDS was also performed using the Bray-Curtis distance index on the occurrences of each species per transect. Finally, diversity between sites was calculated through beta diversity indices thanks to the betapart package (Baselga & Orme, 2012) and its abundance update (Baselga, 2017). First, beta diversity (β_JAC_) was calculated for each transect by transforming the data into presence absence and by using the Jaccard distance index (Jaccard, 1912). Both components of beta diversity were given: species turnover and species nestedness. Then, abundance beta diversity (β_BRAY_) was calculated for each transect by using the Bray-Curtis distance index for abundance data.

## 3 RESULTS

### 3.1 Edge effect distance

Overall, 440 occurrences representing 83 species were collected in total from the six transects (10 m, 30 m, 60 m, 100 m, 250 m and 500 m) at the L site. The wood feeders were the most numerous in occurrences (239) compared to soil feeders (201) but not in terms of species with only 18 wood-feeders species versus 65 species. There were four different families not equally represented. The Termitidae (Amitermitinae, Apicotermitinae, Crepititermitinae, Cylindrotermitinae, Nasutitermitinae, Neocapritermitinae, Syntermitinae, and Termitinae) had 363 occurrences and 77 species, the Heterotermitidae 69 occurrences and 3 species, the Rhinotermitidae 6 occurrences and 2 species while the Kalotermitidae only had 2 occurrences and 1 species (Table 1).

**Table 1.**
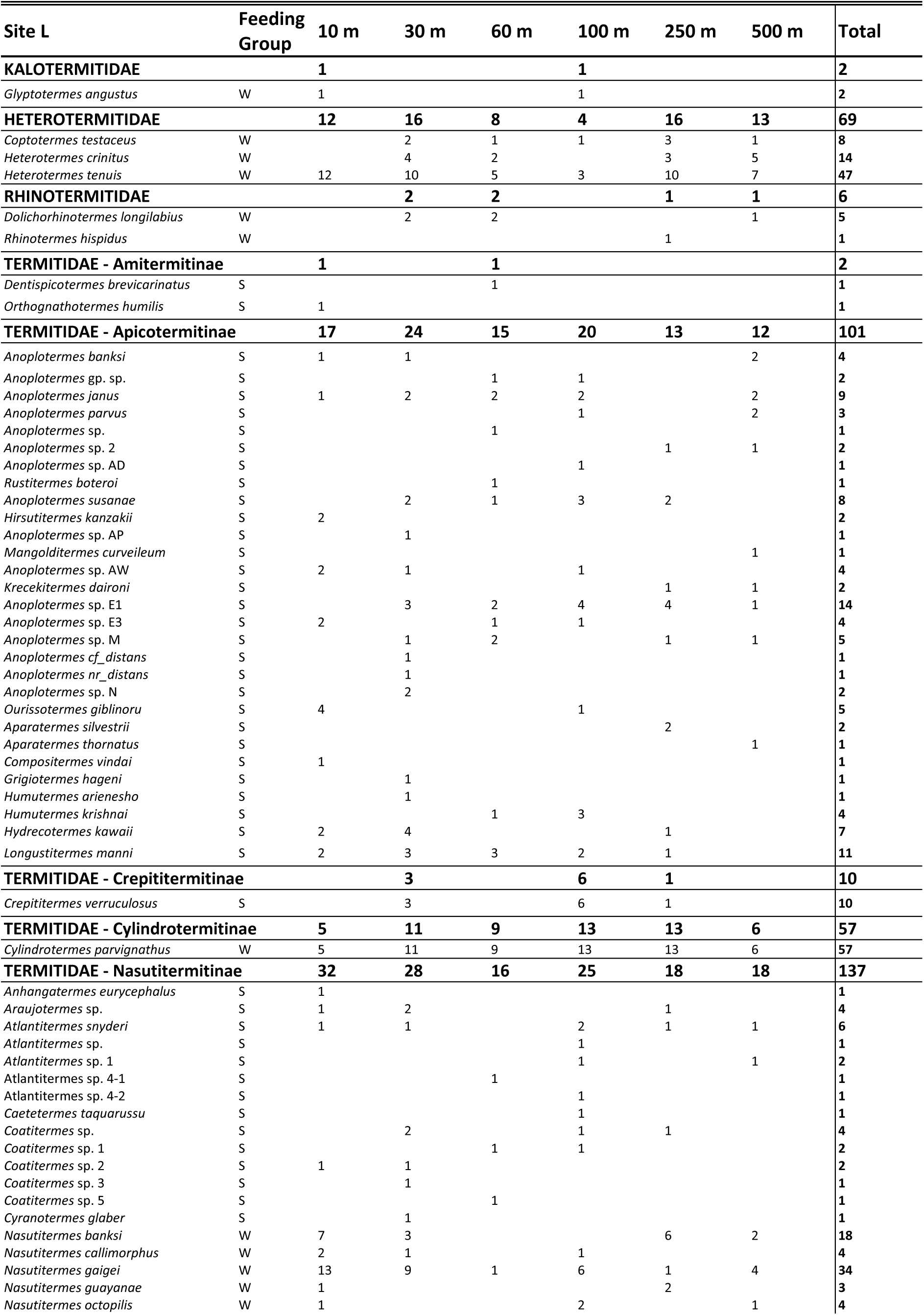
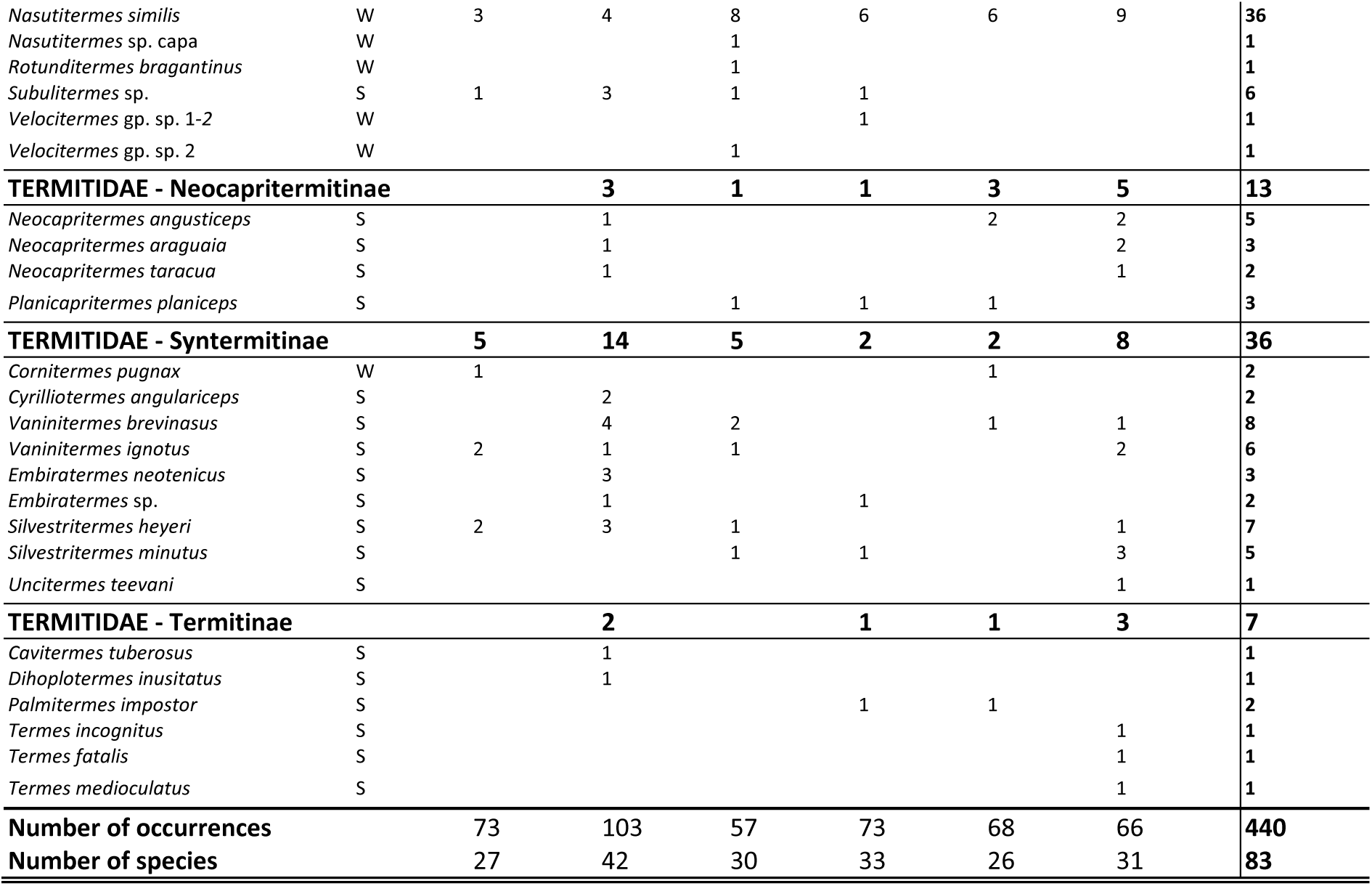
Occurrences of the species (ordered by families and subfamilies) from site L at each transect and in total. Feeding Groups: S = soil feeders and W = wood feeders.

The sample-and occurrence-based rarefaction curves both show that the 30 m and 100 m transects are the richest and the 10 m is the poorest followed by the 250 m transect. There seems to be no consistent pattern between the distance from the edge and the richness. None of the curves reached a plateau, which revealed incomplete sampling (Supplementary S1): completeness, estimated by iNEXT, is high (> 83%) for the first two transects (10 m and 30 m), low (65.8%) for the third transect (60 m) and intermediate (70–78%) for the last three transects (100, 250 and 500 m). Doubling the number of quadrats would increase, on average, the number of species by 39%.

Looking at the environmental variables, elevation seems to get its peak between around 100 m from the forest edge and its lowest at 500 m. The canopy openness (CO) is especially low at 30 m, 250 m and 500 m and the litter mass varies accordingly. Although the CO is the highest at 10 m, the litter mass is not. All variables are significantly different (at α < 0.05) when testing for differences between distances with a Kruskal-Wallis test (elevation: χ^2^ = 133.29, df = 5, p-value < 0.001, CO: χ^2^ = 84.729, df = 5, p-value < 0.001 and Litter: χ^2^ = 48.674, df = 5, p-value < 0.001). There is however no clear pattern between any environmental variable and the distance from the edge. The dead wood volume is highest at 30 m from the forest edge, where it is almost twice higher than along all other transects (Figure 2).

**Figure 2.**
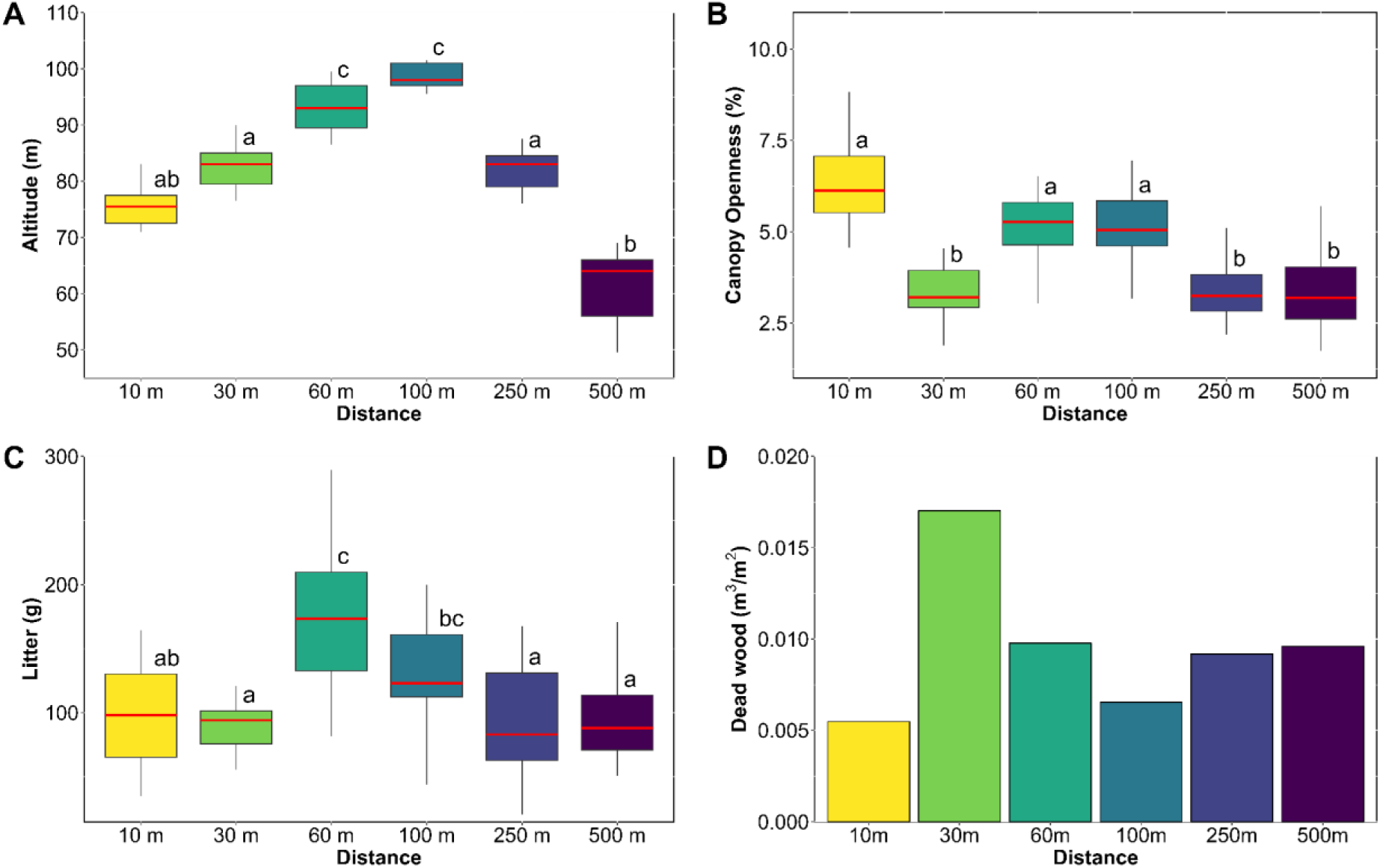
Boxplots of the environmental variables (A = Altitude (m), B = Canopy openness (%) and C = Litter (g)) according to the distance from the forest edge. Two different letters mean that there is a significant difference (α < 0.05) between two distances according to a Dunn post-hoc test (with Bonferroni correction). D is a barplot of the dead wood volume (m^3^/m^2^) for each transect.

Looking at each transect, the 30 m transect was the most diverse with 40 species and 103 occurrences while the 250 m was the least rich with 26 species. The 60 m had the least number of occurrences with 57 occurrences. Looking at the number of species only, the Apicotermitinae were the richest at each transect except at 10 and 100 m where the Nasutitermitinae had slightly more species. The soil feeders were the most diverse for each transect. For the occurrences, the Nasutitermitinae were the most numerous at each site and so were the wood feeders for the transects at 10 m, 60 m, 250 m and 500 m. The soil feeders were the most present at 30 m and at 100 m (Table 1 and Figure 3).

**Figure 3.**
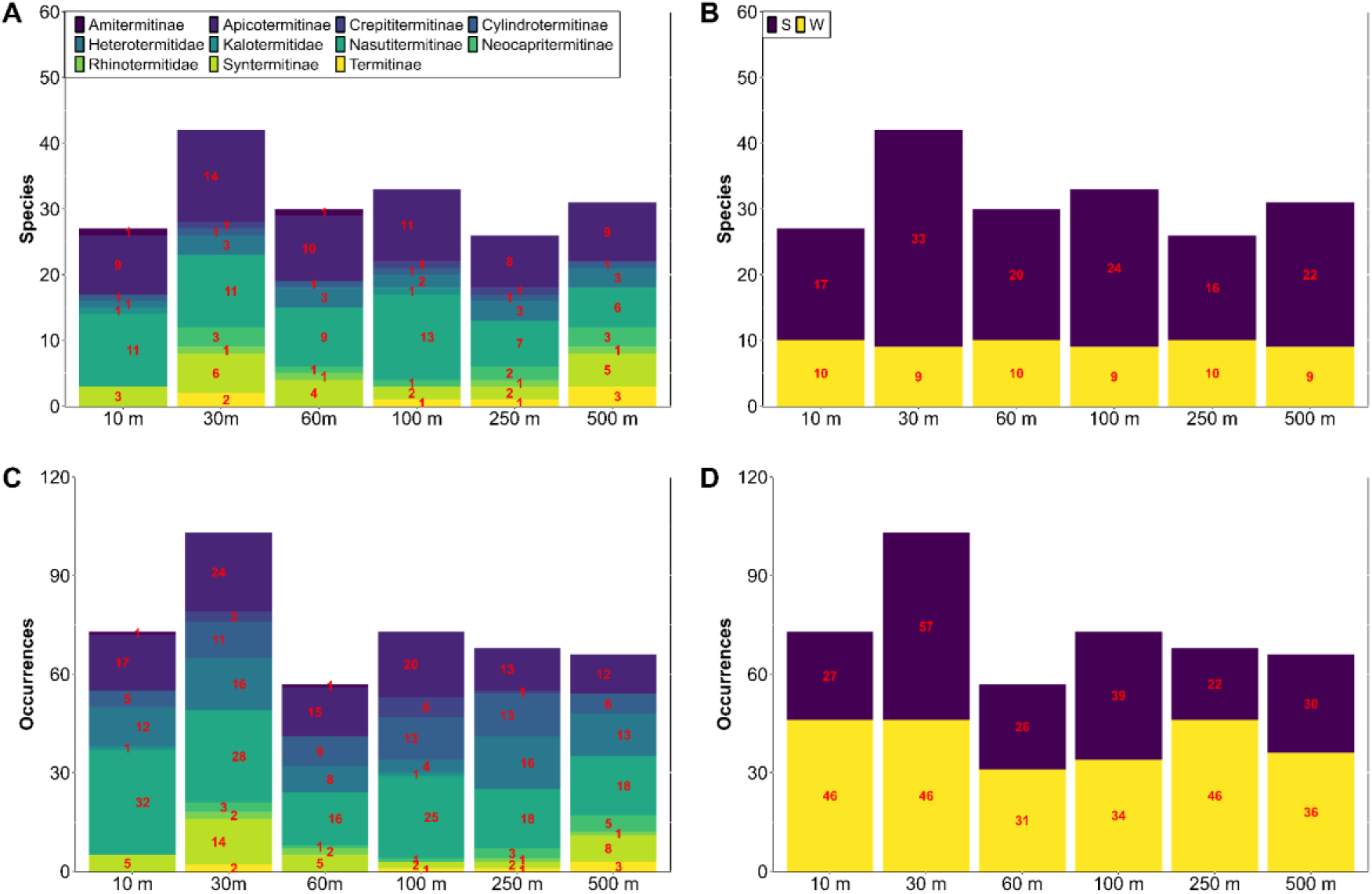
Number of species at each distance from the forest edge (10 m, 30 m, 60 m, 100 m, 250 m and 500 m) for each (sub)-family (A) and each guild (B). Number of occurrences at each distance from the forest edge (10 m, 30 m, 60 m, 100 m, 250 m and 500 m) for each (sub)-family (C) and each guild (D).

No significant differences (α < 0.05) between each transect was found out using a Kruskal-Wallis test of the species relative frequency (χ^2^ = 9.406, df = 5, p-value = 0.094). There weren’t any significant differences as well for each (sub)-family between each transect according to Kruskal-Wallis tests. The wood feeders composition between transects were also not significantly different between each transect (χ^2^ = 0.474, df = 5, p-value = 0.993) and so were the soil feeders (except between the 30 m and the 250 m transect according to a post-hoc Dunn test: χ^2^ = 13.047, df = 5, p-value = 0.023).

The linear regression plots do not show any trend between the Shannon diversity index (H) and altitude nor litter. There seems to be a slight correlation between the canopy closeness and the species diversity; and a good correlation between dead wood volume and species diversity. However, there is no significant correlation between any environmental variable and the Shannon diversity index for all the data or for each feeding guild separately. (Supplementary S2).

The NMDS (Non metric multidimensional scaling) has a good stress of 0.079 and shows that each pair of close by transects (10-30, 60-100 and 250-500) is more similar in terms of communities than more distant transects (Supplementary S3).

### 3.2 Edge effect and beta diversity

Overall, 309 occurrences representing 65 species were collected in total from the six transects at the L, N and M sites. The wood feeders were the most numerous in occurrences (191) compared to soil feeders (118) but not in terms of species with only 19 wood feeders species versus 46 species. There were four different families not equally represented. The Termitidae (Amitermitinae, Apicotermitinae, Crepititermitinae, Cylindrotermitinae, Nasutitermitinae, Neocapritermitinae, Syntermitinae, and Termitinae) had 253 occurrences and 58 species, the Heterotermitidae 48 occurrences and 3 species, the Rhinotermitidae 5 occurrences and 2 species while the Kalotermitidae only had 3 occurrences and 2 species (Table 2).

**Table 2.** Occurrences of the species (ordered by families and subfamilies) from site L (10 m and 100 m), N and M, all the sites at 10 m and at 100 m and in total. Feeding Groups: S = soil feeders and W = wood feeders.

| Sites L, N and M | Feeding group | L10 | L100 | N10 | N100 | M10 | M100 | 10 m | 100 m | Total |
| --- | --- | --- | --- | --- | --- | --- | --- | --- | --- | --- |
| **KALOTERMITIDAE** |  | **1** | **1** |  |  |  | **1** | **1** | **2** | **3** |
| *Glyptotermes angustus* | W | 1 | 1 |  |  |  |  | 1 | 1 | 2 |
| *Kalotermitidae* sp. | W |  |  |  |  |  | 1 |  | 1 | 1 |
| **HETEROTERMITIDAE** |  | **12** | **4** | **16** | **4** | **7** | **5** | **35** | **13** | **48** |
| *Coptotermes testaceus* | W |  | 1 | 1 |  |  |  | 1 | 1 | 2 |
| *Heterotermes crinitus* | W |  |  |  |  |  | 1 |  | 1 | 1 |
| *Heterotermes tenuis* | W | 12 | 3 | 15 | 4 | 7 | 4 | 34 | 11 | 45 |
| **RHINOTERMITIDAE** |  |  |  | **1** | **1** | **1** | **2** | **2** | **3** | **5** |
| *Dolichorhinotermes latilabrum* | W |  |  |  |  |  | 2 |  | 2 | 2 |
| *Dolichorhinotermes longilabius* | W |  |  | 1 | 1 | 1 |  | 2 | 1 | 3 |
| **TERMITIDAE - Amitermitinae** |  | **1** |  |  |  |  |  | **1** |  | **1** |
| *Orthognathotermes humilis* | S | 1 |  |  |  |  |  | 1 |  | 1 |
| **TERMITIDAE - Apicotermatinae** |  | **17** | **20** | **8** | **7** | **6** | **6** | **31** | **33** | **64** |
| *Anoplotermes banksi* | S | 1 |  | 3 | 3 | 2 |  | 6 | 3 | 9 |
| *Anoplotermes* gp. sp. | S |  | 1 |  |  |  |  |  | 1 | 1 |
| *Anoplotermes janus* | S | 1 | 2 |  |  |  | 1 | 1 | 3 | 4 |
| *Anoplotermes parvus* | S |  | 1 |  |  |  |  |  | 1 | 1 |
| *Anoplotermes* sp. AD | S |  | 1 |  |  |  |  |  | 1 | 1 |
| *Anoplotermes susanae* | S |  | 3 |  |  |  |  |  | 3 | 3 |
| *Anoplotermes* sp. AL | S |  |  | 1 |  |  |  | 1 |  | 1 |
| *Hirsutitermes kanzakii* | S | 2 |  | 1 | 1 |  |  | 3 | 1 | 4 |
| *Anoplotermes* sp. AW | S | 2 | 1 |  |  |  |  | 2 | 1 | 3 |
| *Anoplotermes* sp. AX | S |  |  |  |  | 1 |  | 1 |  | 1 |
| *Krecekitermes daironi* | S |  |  |  | 1 | 2 | 1 | 2 | 2 | 4 |
| *Anoplotermes* sp. E1 | S |  | 4 |  |  |  |  |  | 4 | 4 |
| *Anoplotermes* sp. E3 | S | 2 | 1 |  |  |  |  | 2 | 1 | 3 |
| *Anoplotermes* sp. N | S |  |  |  | 1 |  |  |  | 1 | 1 |
| *Anoplotermes* sp. Q | S |  |  | 1 |  |  |  | 1 |  | 1 |
| *Ourissotermes giblinoru* | S | 4 | 1 |  |  | 1 | 1 | 5 | 2 | 7 |
| *Compositermes vindai* | S | 1 |  |  |  |  |  | 1 |  | 1 |
| *Grigiotermes hageni* | S |  |  |  | 1 |  |  |  | 1 | 1 |
| *Humutermes krishnai* | S |  | 3 |  |  |  |  |  | 3 | 3 |
| *Hydrecotermes arienescho* | S |  |  |  |  |  | 2 |  | 2 | 2 |
| *Hydrecotermes kawaii* | S | 2 |  | 1 |  |  | 1 | 3 | 1 | 4 |
| *Longustitermes manni* | S | 2 | 2 | 1 |  |  |  | 3 | 2 | 5 |
| **TERMITIDAE - Crepititermitinae** |  |  | **6** | **1** | **1** | **1** | **1** | **2** | **8** | **10** |
| *Crepititermes verruculosus* | S |  | 6 | 1 | 1 | 1 | 1 | 2 | 8 | 10 |
| **TERMITIDAE - Cyliandrotermatinae** |  | **5** | **13** | **8** | **6** | **5** | **8** | **18** | **27** | **45** |
| *Cyliandrotermes parvignathus* | W | 5 | 13 | 8 | 6 | 5 | 8 | 18 | 27 | 45 |
| **TERMITIDAE - Nasutitermitinae** |  | **32** | **25** | **7** | **21** | **6** | **13** | **45** | **59** | **104** |
| *Anhangatermes eurycephalus* | S | 1 |  |  |  |  |  | 1 |  | 1 |
| *Araujotermes* sp. | S | 1 |  |  |  |  |  | 1 |  | 1 |
| *Atlantitermes oculatissimus* | S |  |  |  |  |  | 1 |  | 1 | 1 |
| *Atlantitermes snyderi* | S | 1 | 2 |  |  |  |  | 1 | 2 | 3 |
| *Atlantitermes* sp. | S |  | 1 |  |  | 1 |  | 1 | 1 | 2 |
| *Atlantitermes* sp. 4-2 | S |  | 1 |  |  |  |  |  | 1 | 1 |
| *Atlantitermes* sp. E3 | S |  | 1 |  |  |  |  |  | 1 | 1 |
| *Caetetermes taquarussu* | S |  | 1 |  |  |  |  |  | 1 | 1 |
| *Coatitermes* sp. | S |  | 1 |  |  |  |  |  | 1 | 1 |
| *Coatitermes* sp. 1 | S |  | 1 |  |  |  |  |  | 1 | 1 |
| *Coatitermes* sp. 2 | S | 1 |  |  |  |  |  | 1 |  | 1 |
| *Nasutitermes banksi* | W | 7 |  |  | 1 |  | 2 | 7 | 3 | 10 |
| *Nasutitermes callimorphus* | W | 2 | 1 | 2 |  |  |  | 4 | 1 | 5 |
| *Nasutitermes gaigei* | W | 13 | 6 | 3 | 4 | 1 | 2 | 17 | 12 | 29 |
| *Nasutitermes guayanae* | W | 1 |  |  | 4 | 1 |  | 2 | 4 | 6 |
| *Nasutitermes octopilis* | W | 1 | 2 |  | 4 | 1 |  | 2 | 6 | 8 |
| *Nasutitermes similis* | W | 3 | 6 | 1 | 6 | 2 | 6 | 6 | 18 | 24 |
| *Nasutitermes* sp. ECU6 | W |  |  |  |  |  | 1 |  | 1 | 1 |
| *Nasutitermes surinamensis* | W |  |  |  | 1 |  |  |  | 1 | 1 |
| *Subulitermes* sp. | S | 1 | 1 | 1 | 1 |  |  | 2 | 2 | 4 |
| *Velocitermes* sp. | W |  |  |  |  |  | 1 |  | 1 | 1 |
| *Velocitermes* gp. sp. 1-2 | W |  | 1 |  |  |  |  |  | 1 | 1 |
| **TERMITIDAE - Neocapritermitinae** |  |  | **1** |  |  |  |  |  | **1** | **1** |
| *Planicapritermes planiceps* | S |  | 1 |  |  |  |  |  | 1 | 1 |

| **TERMITIDAE - Syntermitinae** |  | **5** | **2** | **2** | **10** | **3** | **2** | **10** | **14** | **24** |
| --- | --- | --- | --- | --- | --- | --- | --- | --- | --- | --- |
| *Biratermes robustus* | S |  |  |  |  | 1 |  | 1 |  | 1 |
| *Cornitermes pugnax* | W | 1 |  | 2 | 1 |  |  | 3 | 1 | 4 |
| *Vanitermes ignotus* | S | 2 |  |  | 1 |  |  | 2 | 1 | 3 |
| *Embiratermes* sp. | S |  | 1 |  |  |  |  |  | 1 | 1 |
| *Labiatermes labralis* | S |  |  |  | 1 |  |  |  | 1 | 1 |
| *Silvestritermes heyeri* | S | 2 |  |  | 1 |  |  | 2 | 1 | 3 |
| *Silvestritermes minutus* | S |  | 1 |  | 6 | 2 | 2 | 2 | 9 | 11 |
| **TERMITIDAE - Termitinae** |  |  | **1** | **1** | **2** |  |  | **1** | **3** | **4** |
| *Cavitermes tuberosus* | S |  |  |  | 1 |  |  |  | 1 | 1 |
| *Palmitermes impostor* | S |  | 1 |  |  |  |  |  | 1 | 1 |
| *Termes medioculatus* | S |  |  | 1 | 1 |  |  | 1 | 1 | 2 |
| **Number of occurrences** |  | 73 | 73 | 44 | 52 | 29 | 38 | 146 | 163 | 309 |
| **Number of species** |  | 27 | 33 | 17 | 23 | 15 | 18 | 38 | 56 | 65 |

#### 3.2.1 Comparing diversity between the forest edge and the interior

Both rarefaction curves show that the forest interior is more diverse (species and occurrences) compared to the forest edge. Moreover, the edge almost reaches a plateau which is not the case for the interior (Supplementary S4). Looking more closely with iNEXT, the forest interior has a sample completeness of 78.1% with 57 species while the forest edge has a sample completeness of 89.9% with 38 species. Doubling the number of quadrats could lead to up to 86 species for the forest interior and to 46 species for the forest edge.

Among the occurrences, 146 were from the forest edge (10 m) with 38 species while 163 occurrences were from the forest interior (100 m) with 57 species. The wood feeders were the most numerous at the forest edge (with 97 occurrences and 12 species) and at the forest interior (with 94 occurrences and 19 species). There were 49 occurrences and 26 species of soil feeders at the forest edge while at the forest interior there were 69 occurrences and 38 species. Only one family was more present at the forest edge (35 occurrences and 2 species) than at the forest interior (13 occurrences and 3 species), the Heterotermitidae. The Termitidae had 108 occurrences with 34 species at the forest edge and 145 occurrences with 50 species at the forest interior. The Rhinotermitidae had 2 occurrences with 1 species at theforest edge and 3 occurrences with 2 species at the forest interior while the Kalotermitidae had 1 occurrence at the forest edge and 2 occurrences and species at the forest interior (Table 2 and Figure 4).

**Figure 4.**
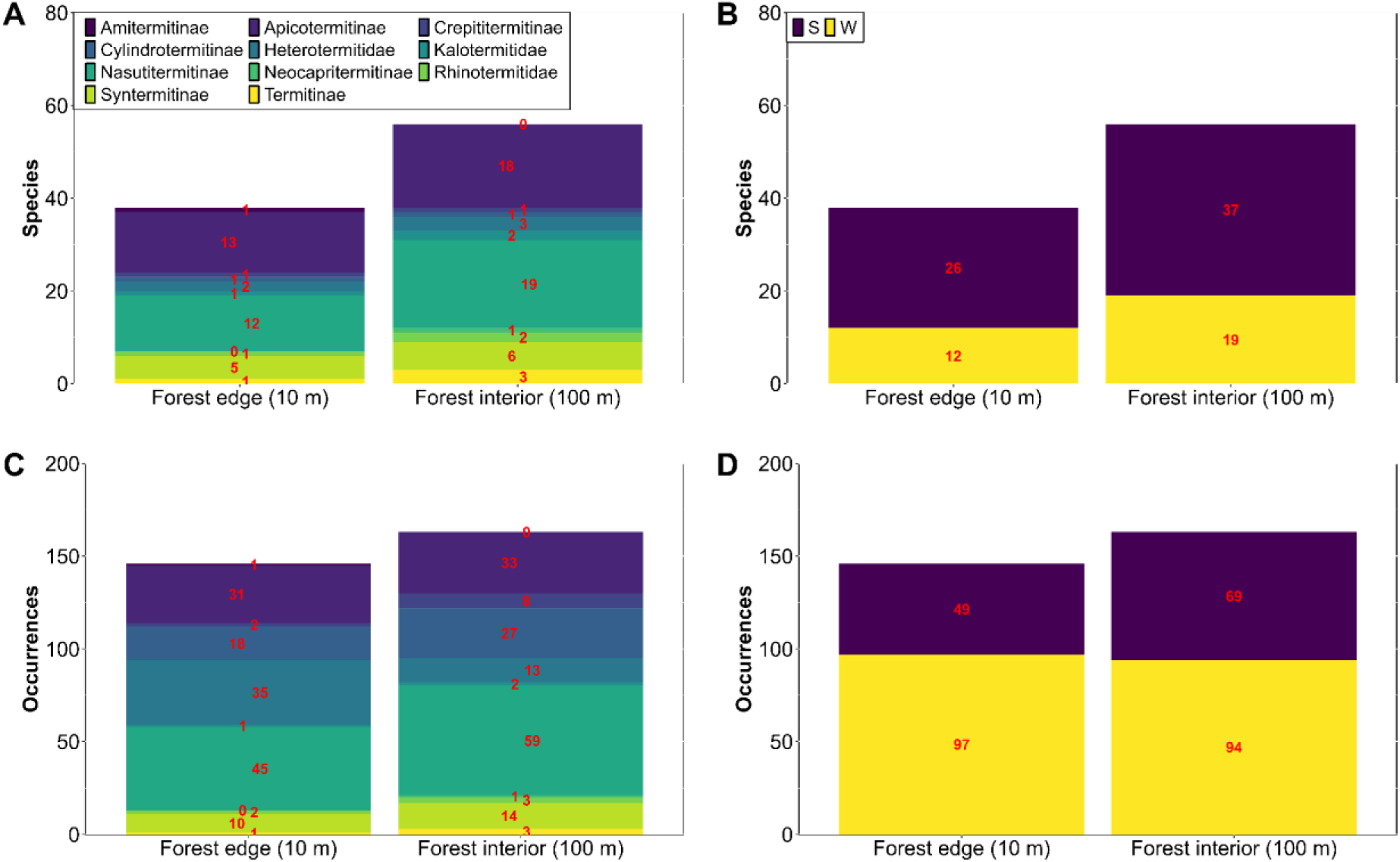
Number of species at the forest edge and at the interior for each (sub)-family (A) and each guild (B). Number of occurrences at the forest edge and at the interior for each (sub)-family (C) and each guild (D).

Overall, the distributions of the forest edge and the forest interior did not significantly differ (α < 0.05) according to a two-sample Kolmogorov-Smirnov test (D = 0.375, p = 0.672). For both guilds, the relative frequency did not significantly differ (α < 0.05) with a Wilcoxon test between the edge and the interior (S: W = 2599, p-value = 0.380; W: W = 2849.5, p-value = 0.884). For the (sub-)families, the relative frequency of the Heterotermitidae differed significantly with a Wilcoxon test between the edge and the interior (W = 3627, p-value < 0.001). When looking at species with enough occurrences to perform Chi-square or Fisher tests (Supplementary S5), only two species have significant differences (α < 0.05) between the forest edge and the interior. *Heterotermes tenuis* (Hagen, 1858) is three times more present at the forest edge (34 occurrences at the edge versus 11 occurrences at the interior; χ^2^ = 16.794, df = 1, p-value < 0.001) and *Nasutitermes similis* (Emerson, 1935) is three times more present at the forest interior (6 occurrences at the edge versus 18 occurrences at the interior; χ^2^ = 7.143, df = 1, p-value = 0.008).

The NMDS has an excellent stress of 0.048 but it does not show any distinction between the forest edge and the forest interior, rather it separates well each site (Supplementary S6).

#### 3.2.2 Comparing diversity between sites

Most species and occurrences were found in the L site with 47 species and 146 occurrences. It is then followed by the N site with 29 species and 96 occurrences and finally by the M site with 25 species and 67 occurrences. Beta diversity between the sites was almost equal between the forest interior (β_JAC_ = 0.74 and β_BRAY_ = 0.56) and the forest edge (β_JAC_ = 0.77 and β_BRAY_ = 0.65). At both distances, the main component was largely species turnover, accounting for more than 90% of beta diversity, not species nestedness.

## 4 DISCUSSION

The diversity and abundance of the three families is similar to other studies realized in the same forest with a large dominance of the Termitidae followed by the Heterotermitidae, the Rhinotermitidae and the Kalotermitidae (Bourguignon *et al*., 2011). Although the Apicotermitinae, with the Nasutitermitinae, is clearly the most diverse and abundant of the Termitidae, it is the most understudied subfamily because most species are soldierless, making identification difficult (Bourguignon *et al*., 2009; Romero Arias *et al*., 2021). With only 3 occurrences and 2 species for all three sites, the Kalotermitidae’s paucity on tropical soils is confirmed, the family being thought to be more present in the canopy (Roisin *et al*., 2006).

### 4.1 Edge effect distance

The rarefaction and extrapolation curves continue to increase slightly when doubling the number of quadrats for every transect. This is especially true for the 60 m transect which has a low completeness compared to the other transects.

No environmental variable seems to be correlated to the distance from the forest edge. Altitude at the site is highly variable with the peak being around 100 m from the forest edge. The canopy openness (CO) is the highest at 10 m but it does not decrease gradually as observed in other studies (Dambros *et al*., 2013, p. 201; Foggo *et al*., 2001). The high CO at 10 m was expected as forest edges are characterized by a higher luminosity compared to forest cores (Laurance, 2004). Litter mass usually increases as the distance from the edge increases as well (Delgado *et al*., 2013) but here it is not the case. Finally, dead wood volume is not linked to edge distance as shown in the results and in Davies (2002). Here, the 30 m transect has the highest dead wood volume, almost twice as much as all the other transects.

That same 30 m transect is also the most diverse of all transects in terms of species and occurrences while the 10 m and 250 m have the lowest species number and the 60 m transect has the least occurrences. Those differences are however not significant, showing that overall, termites are quite resilient against forest edges compared to other insects such as dung beetles or Amazonian ants (Carvalho & Vasconcelos, 1999; Ewers & Didham, 2008; Marsh *et al*., 2018). When looking at families and feeding groups separately, there is as well no significance with distance. There is also no correlation between any environmental variable and the Shannon diversity.

Taking this all together, termites do not really appear to be affected by a forest edge in terms of distance when the forest fragment is large enough to truly neglect habitat reduction effects. This could explain the differences between our results and those of Dambros *et al*. (2013) and Almeida *et al*. (2017).

### 4.2 Edge effect and beta diversity

The rarefaction and extrapolation curves show that the sampling effort was decent with a completeness of more than 78% for the forest edge (10 m) and the forest interior (100 m). The forest interior curve could still increase sharply leading to a bigger gap between the two transects in terms of species diversity.

There was a bit more species and more occurrences at the forest interior than at the forest edge but no significant differences were found. Although, the soil-feeding termites were less present (species and occurrences) at the forest edge than at the forest interior; no significant differences were found whether in terms of species or occurrences. The forest edge does not seem to be too much compelling to soil feeders despite their allegedly thinner skeleton and their food regime requiring a soft and moist soil (Davies *et al*., 2012; Eggleton *et al*., 2002; Muvengwi *et al*., 2017). Neotropical soil feeding termites, and especially wood feeders, seem thus resilient enough against forest edges of higher light intensity and temperature which are not as intense as those found in agricultural cleared land as described by (Eggleton *et al*., 2002). Moreover, Woon *et al*. (2019) found out that soil termites had a higher temperature tolerance than expected but a low desiccation tolerance which, considering the high humidity of the region, might explain the soil feeders’ ability to survive.

For the (sub-)families, the Heterotermitidae were significantly more present at the forest edge due to the important presence of *Heterotermes tenuis* at the forest edge and not so much at the forest interior (three times less). This species, with an immense suitable habitat, is a pest native to South and Central America but considered invasive in the Lesser Antilles and Trinidad and Tobago (Duquesne & Fournier, 2025; Evans *et al*., 2013). Moreover, *H. tenuis* is able to exploit several food sources and under different environmental conditions (Arab *et al*., 2005). One hypothesis would be that when the forest was cut in half, *H. tenuis* was able to quickly colonize and dominate the forest edge before other species thanks to its opportunistic abilities. This was already shown for ants where opportunistic species, thanks to their good adaptability and dispersibility, were more present at forest edges than cryptic species (Ivanov & Keiper, 2010). Finally, Caitano *et al*. (2020) has also shown that forest edges have an overall positive effect on invasive insect species. *Nasutitermes similis*, a wood feeder, was three times more present at the forest interior than at the forest edge. However, other *Nasutitermes* species were present equally at the forest edge and at the interior, giving us no plausible explanations for its low presence at the forest edge.

Finally, beta diversity was almost equal for both transects and can be explained by species turnover by 90%, and not by nestedness. These results do not follow those predicted by the landscape-divergence hypothesis from Laurance *et al*. (2007) which leads to higher beta diversity near the edge. Here, the results show that termites are resilient species able to cope with a forest edge, especially wood feeders as soil feeders were a bit less present. Species composition did not significantly differ between the forest edge and the core, on the contrary to Dambros *et al*. (2013) and Almeida *et al*. (2017) findings. For those two studies, the differentiation probably came from the fragments size which were too small to discriminate true edge effects from those of habitat reduction.

## 5 CONCLUSION

The effects of a forest edge, independently from reduced fragment size effect, were lower than expected. Here, neotropical termites were not affected 100-300 m into the interior although several other studies focusing on ecologically close insects found the opposite. Soil feeders were not significantly less present at the forest edge although they have an allegedly thinner skeleton and a need for moist and soft soil. We also did not find any increase in beta diversity near the forest edge notwithstanding the landscape-divergence hypothesis of Laurance *et al*. (2007). Overall, neotropical termites seem to be resilient insects against forest edges as long as the fragment size stays large enough. However, we found a larger presence of an opportunistic and pest species (*Heterotermes tenuis*) at the forest edge. Its good adaptability and dispersibility may have allowed the species to quickly colonize this new environment before more cryptic species, as it’s been shown for ants. As deforestation and forest edges increase in tropical forests, we expect opportunistic and potentially invasive termites to be more present globally. Finally, we suggest that future studies take more into account fragment sizes when studying the impact of forest edges to truly discriminate its effects.

## Supporting information

DarwinCore dataset

## ACKNOWLEDGMENTS

This paper is entirely dedicated to the loving memory of Arthur Renard who would have been an exceptional tropical entomologist and would undoubtedly have contributed to this research. Furthermore, this work is dedicated to our fellow co-author and friend, Yves Roisin, whose dedication to the termite world will never be forgotten. This research was funded by the French Community of Belgium through a grant from the FNRS (J0193-25) to Yves Roisin and from FNRS/FRIA funds to Edouard Duquesne. We would also like to thank Nicolas Fontaine for his help with the genetic analyses and Loane Wu for her help on the field and in the lab.

## AUTHOR CONTRIBUTION

Edouard Duquesne: Data curation, Formal Analysis, Investigation, Methodology, Visualization, Writing – original draft, Writing – review & editing. Antoine Dumortier: Data curation, Investigation.

Camille Legrand: Data curation, Investigation.

Pavel Babczenko: Data curation, Investigation.

Yves Roisin: Conceptualization, Funding acquisition, Investigation, Methodology, Project administration, Resources, Supervision, Writing – review & editing.

