## Supporting information for "Forest edge effects on termites in a neotropical rainforest"

**Supplementary S1.** Sample- (left) and occurrence-based (right) rarefaction curves for each distance from the forest edge.

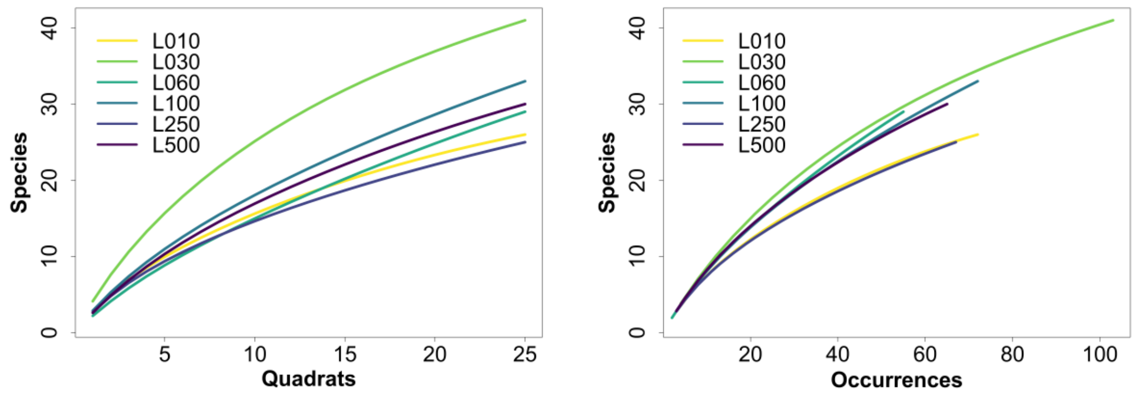


**Supplementary S2.** Linear regression plots between the Shannon diversity index of each distance and between an environment variable (A = Altitude (m), B = Canopy openness (%), C = Litter (g) and D = Dead Wood (m3/m2)). R2 adjusted values and p-value are added for each plot.

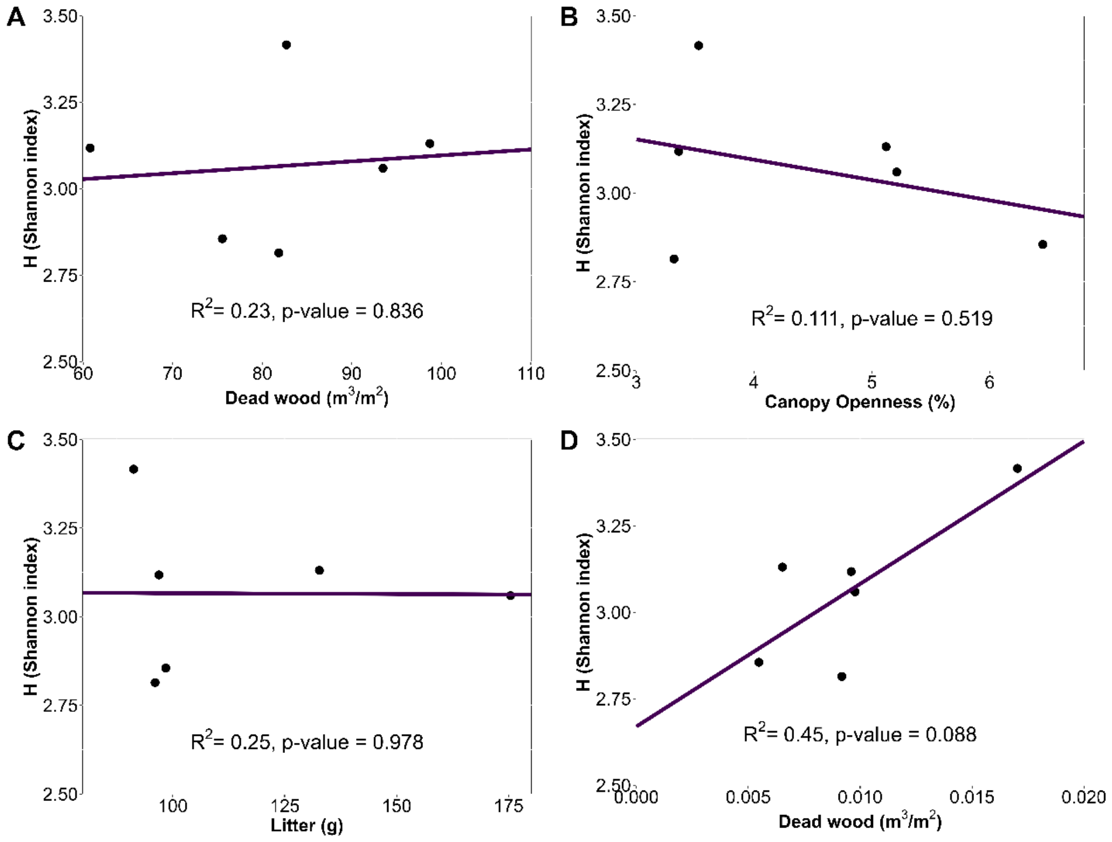

**Supplementary S3.** Non-metric multidimensional scaling plot. Points represent transects at a distance from the edge. The closer sites are, the more similar to one another they are.

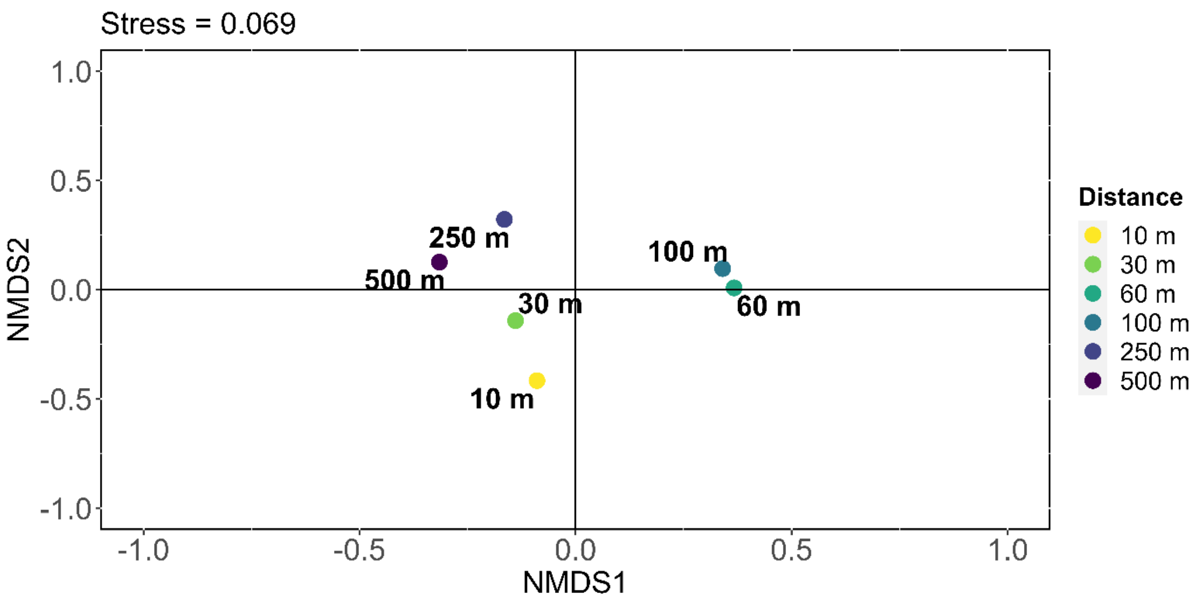


**Supplementary S4.** Sample- (left) and occurrence-based (right) rarefaction curves for the forest edge (10 m) and the forest interior (100 m).

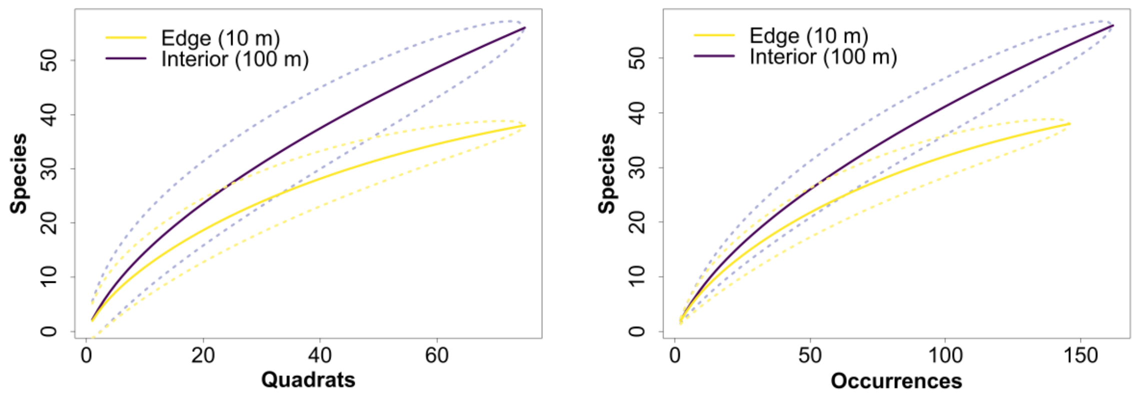

**Supplementary S5.** Values of Chi-square and Fisher tests for species with enough occurrences (see Methods). Species in bold have a significant difference between the forest edge and the forest interior (∝ < 0.05).

| Chi-Square | 10 m | 100 m | χ2 | df | p-value |
|---|---|---|---|---|---|
| *Nasutitermes gaigei* | 17 | 12 | 1.069 | 1 | 0.301 |
| ***Nasutitermes similis*** | 6 | 18 | 7.143 | 1 | **0.008** |
| *Cylindrotermes parvignathus* | 18 | 27 | 2.571 | 1 | 0.109 |
| ***Heterotermes tenuis*** | 34 | 11 | 16.794 | 1 | **<0.001** |
| **Fisher** | **10 m** | **100 m** | | **p-value** | |
| *Anoplotermes banksi* | 6 | 3 | | 0.494 | |
| *Anoplotermes janus* | 1 | 3 | | 0.620 | |
| *Anoplotermes susanae* | 0 | 3 | | 0.245 | |
| *Hirsutitermes kanzakii* | 3 | 1 | | 0.620 | |
| *Anoplotermes sp. AW* | 2 | 1 | | 1.000 | |
| *Krecekitermes daironi* | 2 | 2 | | 1.000 | |
| *Anoplotermes sp. E1* | 0 | 4 | | 0.120 | |
| *Anoplotermes sp. E3* | 2 | 1 | | 1.000 | |
| *Ourissotermes giblinoru* | 5 | 2 | | 0.442 | |
| *Hydrecotermes kawaii* | 3 | 1 | | 0.620 | |
| *Longustitermes manni* | 3 | 2 | | 1.000 | |
| *Atlantitermes snyderi* | 1 | 2 | | 1.000 | |
| *Nasutitermes banksi* | 7 | 3 | | 0.327 | |
| *Nasutitermes callimorphus* | 4 | 1 | | 0.367 | |
| *Nasutitermes guayanae* | 2 | 4 | | 0.681 | |
| *Nasutitermes octopilis* | 2 | 6 | | 0.276 | |
| *Subulitermes sp.* | 2 | 2 | | 1.000 | |
| *Cornitermes pugnax* | 3 | 1 | | 0.620 | |
| *Vaninitermes ignotus* | 2 | 1 | | 1.000 | |
| *Silvestritermes heyeri* | 2 | 1 | | 1.000 | |
| *Silvestritermes minutus* | 2 | 9 | | 0.056 | |
| *Crepititermes verruculosus* | 2 | 8 | | 0.098 | |
| *Dolichorhinotermes longilabius* | 2 | 1 | | 1.000 | |

**Supplementary S6.** Non-metric multidimensional scaling plot. Blue points are sites at the forest edge (10 m) while red points are sites at the forest interior (100 m). The closer sites are, the more similar to one another they are.

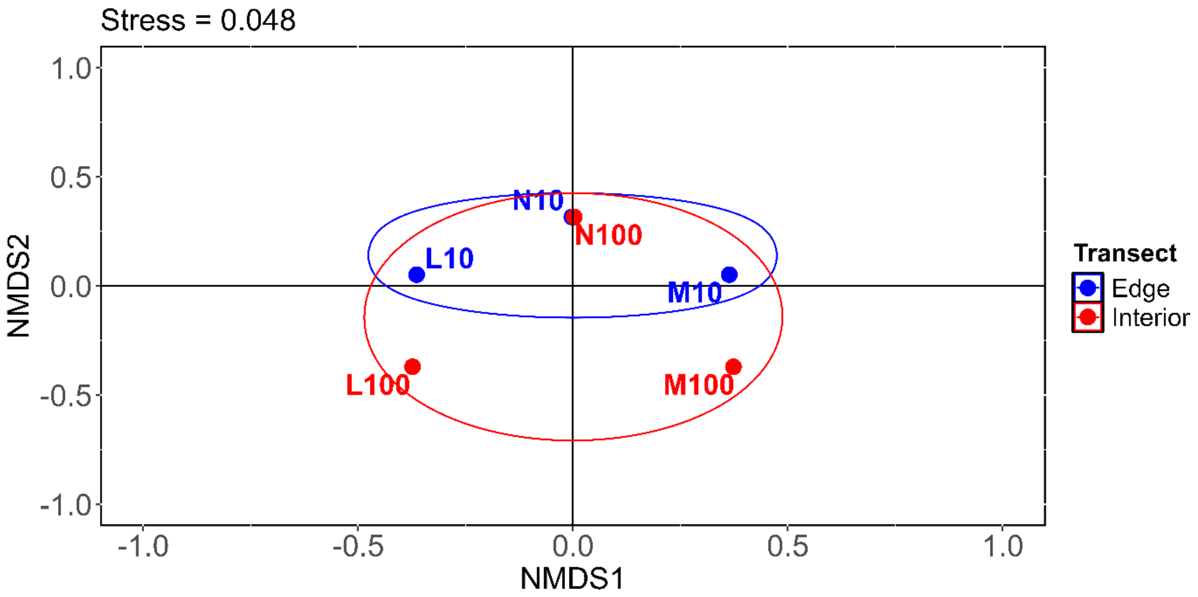
